# Arterial Shear Rate Determines the Structure and Mechanical Properties of Whole-Blood Clots

**DOI:** 10.64898/2025.12.05.692588

**Authors:** Hande Eyisoylu, Parvin Ebrahimi, Marla Lavrijsen, Moniek P.M. de Maat, Gijsje H. Koenderink

## Abstract

**Background:** Arterial thrombosis arises under high shear conditions and is a leading cause of ischemic stroke and myocardial infarction. The local hemodynamic environment during the formation of the thrombus influences its final composition, which in turn influences its mechanical characteristics and the associated risk of embolization. However, how arterial shear rates determine the local mechanical properties, structure and composition of clots remains poorly understood.

**Objective:** To investigate how physiological and pathological arterial shear rates shape blood clot composition, formation dynamics, and regional mechanical properties in a newly-developed microfluidic flow model.

**Methods:** We used a microfluidic flow model that allows real-time imaging of blood clot formation under controlled arterial shear rates (300, 1000, and 3500⍰s^−1^). We visualized platelets and fibrin deposition using confocal microscopy and quantified composition-dependent spatial heterogeneities in clot stiffness with microindentation.

**Results:** Pathologically high (3500 s^-1^) arterial shear rates promoted more rapid platelet aggregation and the formation of taller platelet-rich clots. Fibrin deposition started after about 6 minutes in all shear conditions and coincided with platelet aggregate stabilization at higher shear rates. Microindentation revealed that platelet- and fibrin-rich regions were significantly stiffer than red blood cell-rich regions. Clots formed at 3500⍰s^−1^ had a > 3-fold higher average stiffness than those formed at 300⍰s^−1^, reflecting their higher platelet content.

**Conclusions:** This study demonstrates that arterial shear rate governs clot growth and composition, translating into shear-dependent microscale mechanical properties. These findings provide new mechanistic insights into clot properties that are relevant for clot stability and vulnerability to embolization.

## Introduction

Arterial thrombosis is a critical underlying cause of cardiovascular events, such as myocardial infarction and ischemic stroke [1]. In arteries, thrombi arise under a high flow, often on ruptured atherosclerotic plaques that trigger thrombus formation via the exposure to thrombogenic materials like tissue factor, collagen, and Von Willebrand Factor (VWF) [1]. The local hemodynamic environment influences the initiation, propagation, and ultimate physical characteristics of the arterial thrombus. As blood flows through the arteries, variations in vessel diameter and flow velocity generate gradients in the vessel. Accordingly, within the arterial tree, shear rates can vary significantly, ranging from relatively low arterial shear (∼300 s^−1^ in larger arteries) to moderate (∼1000 s^−1^ in arterioles) and pathological shear rates (>3000 s^−1^ in stenosed vessels), while extremely high shear rates may reach beyond 10,000 s^−1^ in severely stenotic cases [2, 3]. This wide spectrum of arterial shear rates influences thrombus structure and composition, which will in turn influence mechanical properties of the thrombus [4].

Mechanical properties of arterial thrombi such as elastic modulus and fracture strength are major determinants of their mechanical stability, influencing the likelihood of embolization and the potential to successfully remove clots by mechanical thrombectomy [5]. Red blood cell (RBC)-rich thrombi are easier to remove via thrombectomy compared to fibrin-rich thrombi, but fibrin-rich thrombi are less likely to fragment during this procedure [6-8]. Similarly, stroke thrombi that are rich in RBCs lyse more easily during intravenous thrombolysis [6, 9]. Blood composition, platelet reactivity, and arterial geometry, thus local shear and flow, vary between individuals. This complex interplay of different factors that influence clot properties obscures the specific contribution of flow to its composition, structure, and mechanics. This complexity motivates the use of *in vitro* models in which individual variables can be isolated and systematically tested under controlled conditions.

*In vitro* microfluidic models have become useful tools for studying thrombosis under flow conditions using small samples of human blood and providing controlled shear environments that approximate *in vivo* conditions. *In vitro* flow studies have shown that the shear rate determines the role of VWF in thrombus formation. At high arterial shear, VWF unfolds and binds platelet GPIbα receptors to drive rapid tethering and aggregation, with subsequent stabilization via the integrin receptor GPαIIbβ3 that binds fibrin [10-12]. At lower shear rates, thrombus formation is more coagulation-driven, producing fibrin- and RBC-rich thrombi with fewer platelets [13-15]. Finally, several studies reported an increased occlusion capacity of thrombi with increasing shear rate [16, 17].

However, a knowledge gap remains regarding how arterial shear rates influence the spatial heterogeneity of a thrombus’s mechanical properties. Most mechanical measurements of *in vitro* clots until now have been performed at a macroscopic scale [18]. With few exceptions [14, 19], mechanical measurements have been restricted to clots formed under static conditions. These readouts are global, while stroke clots are known to exhibit marked spatial heterogeneity with distinct RBC-rich vs. platelet/fibrin-rich regions [20, 21]. It is crucial to characterize the mechanical heterogeneities associated with this compositional heterogeneity for accurate predictions of *in vivo* embolization risk or thrombectomy outcomes. Therefore, it is necessary to link the dynamic processes of thrombus formation under arterial shear conditions to their effects on the local mechanical stiffness of the developing clot.

For this study, we developed a microfluidic thrombus-on-a-chip model that allows precise control over the shear rate during clotting, mimicking physiologically relevant and pathological flow conditions. Using this platform, we investigated thrombus formation under arterial shear rates spanning from physiological (300 s^−1^, 1000 s^−1^) to pathological levels (3500⍰s^−1^), combining real-time confocal imaging of platelet and fibrin deposition with post-clotting microindentation to map regional mechanical properties. By linking the arterial shear history of clot formation to the detailed composition, structure, and local mechanics, our work may help advance understanding and prediction of clinical outcomes such as clot stability and response to thrombectomy in arterial thrombosis.

## Methods

### Microfluidic Model

#### Device Design

The thrombus-on-a-chip model was designed to mimic *in vivo* flow conditions while mitigating the risks of premature embolization or flushing of the clot due to increased pressure buildup during clot growth. Our device includes a single inlet and two outlets inspired by Berry et al [22]. The two parallel channels have rectangular cross-sections with dimensions of 500⍰μm in width and 70⍰μm in height (SI Figure 1), ensuring a height-to-width ratio that allows for platelet aggregation [23]. Additionally, the device contains a vacuum chamber that enables reversible bonding of the PDMS chip to the glass coverslip; upon release of the vacuum, the channel can be opened to directly access the formed clot for mechanical characterization. A detailed description of the device fabrication and assembly can be found in the Supplemental Information.

#### Surface Coating with Collagen and Tissue Factor

To simulate physiological thrombus formation, a thrombogenic spot was created on the surface of a glass coverslip (VWR, #1) by coating with collagen I and tissue factor [24]. Briefly, a 2 μL droplet of Collagen I Horm suspension (Takeda; 0.1 mg mL) was pipetted onto the coverslip and left to dry for one hour in a humid environment at room temperature. Next, a 2 μL droplet of tissue factor (Siemens, 200 pM) was pipetted on the same spot and left to dry for another hour. To make sure the coated spot was consistent in size (1000 x 500 μm), a mask was taped over the glass coverslip. The coated spot was positioned centrally in one of the parallel arms of the device to direct clot formation to the desired region.

### Experimental Setup

#### Blood collection and preparation

Blood was collected via venipuncture from 4 healthy volunteers into 9 ml 3.2% sodium citrate tubes (Vacuette, Greiner Bio-one). Approval for the study was obtained from the local Medical Ethics Committee (MEC-2021-0849). All experiments were conducted within 4 hours of the blood collection. Blood was labeled with DiOC6 (final concentration 0.5 µg/mL, Anaspec, Reeuwijk, The Netherlands) for platelets and with Alexafluor AF647 fibrinogen (final concentration 16.5 µg/mL, Molecular Probes, Life Technologies, New York NY, USA) for fibrin, and with fluorescently labelled (AF594) VWF antibody (Dako, 1:400). Immediately before blood was perfused through the channel, it was recalcified with CaCl_2_ (final concentration 17mM). Experiments using human blood were in accordance with established guidelines for thrombosis research [23, 25, 26].

#### Realtime Imaging of Clot Formation

Clots were formed under three different arterial shear rates (300 s^-1^, 1000 s^-1^, 3500s^-1^) controlled using a syringe pump (NE-1000 One Channel Programmable Syringe Pump). The wall shear rates γ (s^-1^) were computed from the volumetric flow rate *Q* (ml/min) using the Poiseuille solution for a rectangular duct [25]:

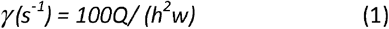

Due to the Y-shaped channel with two identical parallel arms, the volumetric flow rate was doubled in the syringe pump settings to achieve the desired wall shear rates. Blood was perfused through the channels and over the coated patch to initiate clotting. Flow was maintained for 10 minutes.

Real-time imaging of clot formation was performed using a Leica TCS SP8 confocal microscope (Leica Microsystems) equipped with an HC PL APO CS2 20×/0.75 NA dry objective. To visualize platelet and fibrin deposition, 488 nm and 633 nm lasers were used, respectively. Image acquisition was conducted in a time-lapse z-stack mode with a scanning resolution of 512 × 512 pixels and a scan speed of 600 Hz.

To capture the entire height of the microchannel and the developing clot, z-stacks spanning 70⍰µm in depth were acquired at a step size of 2⍰µm. Time-lapse imaging was performed over a 10-minute period, with full z-stacks captured every 16 seconds. To ensure complete spatial coverage of the coated region, two regions of interest (ROIs) were defined across the flow chamber based on the markings indicating the coated region. The confocal system was programmed to sequentially acquire z-stacks at ROI 1 and ROI 2 in alternating fashion throughout the imaging period, enabling near-continuous visualization of thrombus growth over time.

#### Image Analysis

All image processing and quantification were performed using Fiji (ImageJ, NIH)[27]. Confocal z-stack time series acquired during clot formation were converted into maximum intensity projections for each time point (1–10 minutes). To isolate clot regions in the platelet and fibrin channels, automated thresholding was applied separately to each channel. After visual inspection of several thresholding methods (see SI Figure 2), the Triangle algorithm was selected for the platelet channel, and the Otsu method was used for the fibrin channel, as these provided the most accurate segmentation of the clot structure in their respective channels. Following thresholding, we generated binary masks and quantified the percentage of the surface area covered by platelets and fibrin for each time point. From this analysis, we determined the kinetics of clot growth and the final clot composition. The clot height was quantified using the full z-stack at the final time point. The maximum height was defined as the vertical extent of continuous clot signal based on either channel. To assess the average lateral size of platelet aggregates, maximum intensity projections of the platelet channel at the final time point were used. Following thresholding and mask generation, we calculated the average size of individual platelet aggregates with the “Analyze Particles” tool in Fiji.

#### Mechanical Characterization

After the clots were formed under either the lowest (300 s^−1^) or highest (3500 s^−1^) shear rate, the vacuum bond was released and the PDMS channel was carefully separated from the glass coverslip. The coverslips with the thrombi were submerged in HEPES (4-(2-hydroxyethyl)-1-piperazineethanesulfonic acid) buffer (10 mM HEPES, 115 mM NaCl, pH 7.4) to preserve the clot structure. To assess the local mechanical properties of the thrombi, microindentation was performed using a Chiaro Nanoindenter (Optics11 B.V., Amsterdam, NL) operated by the Piuma software (Optics11 Life). The indenter was mounted on an epifluorescence microscope (Leica Thunder) equipped with a 10x dry objective (HC PL APO 0.45NA, Leica) and monochrome sCMOS camera (Leica). The combined setup enabled us to correlate local indentation measurements with the local clot composition directly at the site of indentation. Clots were fluorescently labeled for platelets and fibrin (same as for confocal imaging) and imaged using a 200 mW solid state LED5 light source (Leica), while RBCs were visualized via the brightfield channel. This visual guidance allowed us to classify indentation sites as “fibrin and platelet rich”, “RBC and fibrin rich,” or “RBC rich” regions. Specifically, for each indentation region, we checked each channel (platelet (fluorescence), fibrin (fluorescence) and RBCs (brightfield)) for a signal. If there was a signal from any of these channels the region was classified as rich in that component (for examples, see Fig. 4D).

Indentations were performed in indentation control mode using a spherical probe with a radius of 56 µm and stiffness of 0.22 N/m. Each indentation was performed at a loading and unloading rate of 2 µm/s, to an indentation depth of 2 µm, making sure that the maximum indentation depth was less than 5-10% of the sample thickness to avoid any influence of the substrate on the stiffness measurements.

The resulting force (*F*) - indentation (*δ*) curves were analyzed using the Hertzian contact model to calculate the Effective Young’s modulus (E_eff_). Specifically, the loading branch was fitted to the Hertz model for a rigid sphere indenting an elastic half-space [28]:

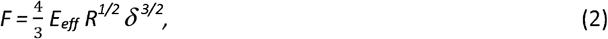

The effective Young’s modulus is related to the actual Young’s modulus *E* according to:

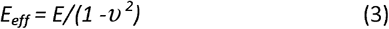

We assumed a Poisson’s ratio (*υ*) of 0.5, corresponding to an incompressible material. Curve fits were performed with the Optics11 Data Viewer software, which implements a least-squares fit of the measured *F–δ* data to Equation 2 (see SI Figure 3 for example). Because clots are viscoelastic and heterogeneous, we interpret E_eff_ as a comparative local stiffness rather than an absolute elastic modulus.

### Statistical Analysis

All image-based quantifications were performed using clots generated from four healthy donors, with four (total n=16) clots analyzed per donor for each shear condition (300 s^−1^, 1000 s^−1^, and 3500⍰s^−1^). For mechanical characterization, microindentation testing was performed on a total of 9 clots for both the 300 s^−1^ and 3500⍰s^−1^ conditions, with at least 5 indentations conducted per clot in compositionally distinct regions identified by confocal imaging. Data are presented as mean ± standard error. Statistical comparisons between shear conditions were performed using one-way ANOVA with Tukey’s post hoc test for multiple comparisons. For non-parametric data, the Kruskal-Wallis test with Dunn’s correction was applied. A p-value < 0.05 was considered statistically significant. All statistical analyses were conducted using GraphPad Prism (version 10, GraphPad Software).

## Results

### Real-Time Observation of Clot Formation

We visualized clot formation in real time with confocal microscopy under conditions of arterial flow using a microfluidic flow model. Blood was flown into the device at different flow rates to achieve shear rates representing low (300 s^-1^) and mid (1000 s^-1^) normal arterial flow rates and a pathologically high flow rate (3500s^-1^). Figure 1 shows representative confocal images of platelets and fibrin over time, comparing the clotting dynamics at the three shear rates. To quantify the temporal progression of platelet accumulation and fibrin deposition, we determined the percentage of surface area covered by platelets and fibrin, illustrated with representative examples underneath the curves. Representative movies showing 3D renderings of platelet, fibrin and merged channels over time for each shear rate are presented in Supplemental Movie 1.

**Figure 1:**
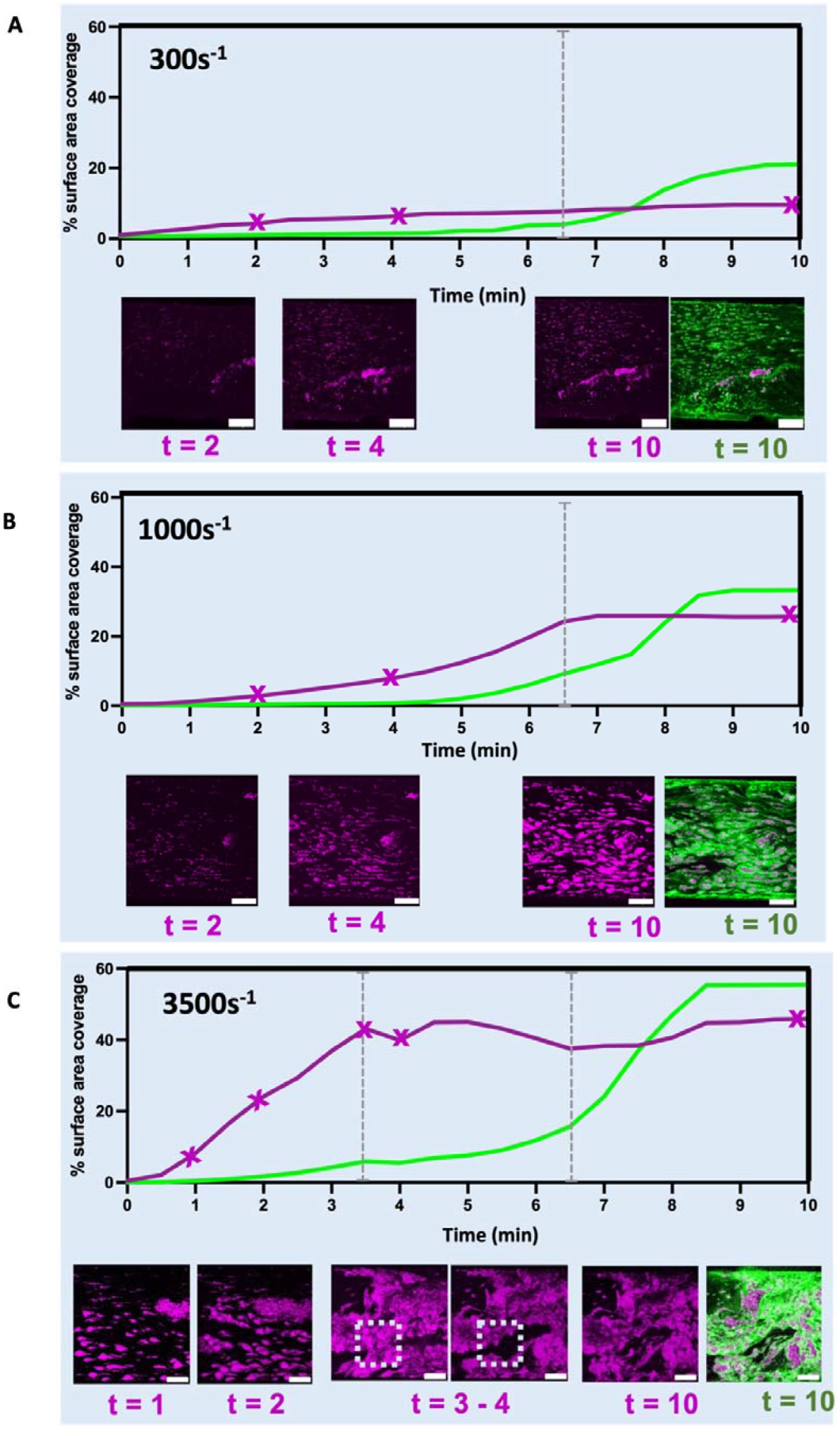
Temporal dynamics of platelet deposition and fibrin network formation under arterial shear conditions. Representative examples of the evolution of the fibrin (green) and platelet (magenta) surface coverage (in %) over a period of 10 minutes at shear rates of (A) 300s^−1^, (B) 1000 s^−1^, and (C) 3500⍰s^−1^. Underneath the plots, maximum intensity confocal projections of platelets at early, mid, and late time points are shown (the corresponding timepoints (minutes) are indicated by the x signs on the magenta curve for each image) along with a merged maximum intensity projection of the final clot. The mid (3-4 min) timepoint is shown only for the 3500s^-1^, to highlight the unstable platelet aggregation during this period. For the lower 300s^-1^ and 1000s^-1^ shear rates the early and the late timepoints are shown. (A) At 300s^−1^, platelet adhesion begins within the first 2 minutes, forming small, discrete aggregates. Platelet surface coverage remains low throughout the 10-minute period. Fibrin network formation begins around 6 minutes (indicated by the dashed line). (B) At 10001s^−1^, platelet surface coverage gradually increases, reaching a plateau after 6 minutes, coinciding with the onset of fibrin network formation (indicated by the dashed line). (C) At 3500 s^−1^, the platelets show rapid aggregation within the first 3 minutes, followed by a transient unstable phase (between the two dashed lines) that ends after 6 minutes, as the fibrin network starts to form and stabilize the clot. The confocal images show that during the unstable phase, platelets that were attached in the earlier time frame (shown inside the dashed square) are washed away in the next frame. Scale bars: 100µm.

At the lowest shear rate (300⍰s^−1^, Fig. 1A), platelet adhesion to the bottom of the microfluidic channel began within the first 2 minutes, forming small, discrete aggregates. Platelet surface coverage remained low throughout the 10-minute observation period. The platelet aggregates themselves also remained unchanged, showing no evidence of rearrangement or fragmentation. Fibrin deposition started after 6 minutes, as indicated by the dashed line (Fig. 1A), typically around the platelet aggregates (Supplementary Figure 4). At the shear rate of 1000⍰s^−1^, platelet aggregates formed across the entire coated surface and were significantly larger than those observed at 300⍰s^−1^ (Fig. 1B). The platelet coverage gradually increased, reaching a plateau after 6 minutes. Fibrin deposition again began after 6 minutes, localizing around platelet clusters. At the highest shear rate of 3500⍰s^−1^, platelet adhesion and aggregation was much more rapid, with coverage rising sharply to ∼40% within 3 minutes (Fig. ⍰1C). However, following the rapid aggregation phase, platelet aggregates were unstable, showing flow-induced rearrangement and partial detachment (Fig. 1C). This unstable behavior persisted until approximately 6 minutes, after which the coverage of platelets stabilized. This stabilization moment coincided with the onset of fibrin deposition, which again started around 6 minutes. Fibrin deposition originated at platelet aggregates and gradually extended to interconnect the aggregates.

Additionally, using the brightfield channel, we visualized RBC dynamics relative to platelet and fibrin architecture in real time. Supplemental movies 4-6 of the brighfield channel show RBC entrapment over time. Following initial platelet adhesion and aggregation, RBCs began to colocalize with the platelet aggregates. As a fibrin network developed around these platelet aggregates and some RBCs, RBC retention became more stable while free floating RBCs were still able to flow around. For all shear rates, an initial platelet adhesion and aggregation over the coated surface was followed by RBC entrapment via a final fibrin network formation.

In summary, larger shear rates promote faster and more extensive platelet aggregation than lower shear rates. Across all shear rates, fibrin formation started after 6 minutes and was spatially organized around platelet aggregates, which acted as nucleation points for fibrin fiber extension and clot stabilization (Supplementary Figure 4).

### Shear-dependent platelet recruitment is potentially mediated by VWF

The confocal data reveal a striking and rapid enhancement of platelet accumulation at 3500 s^−1^. A likely candidate mechanism is shear-sensitive unfolding of VWF [29]. To test this mechanism, we performed flow experiments in the presence of a fluorescently tagged antibody specific for VWF. At the high (3500 s^-1^) shear rate, we observed clear colocalization of the platelet aggregates with VWF (Figure 2C). Long strands of VWF elongated in the flow direction already appeared within a minute after initiating clotting (Supplemental Movie 2 of VWF over time). By contrast, clots formed under lower shear rates (300 s^-1^ or 1000 s^-1^) showed little VWF signal and no evidence of VWF strands (Fig. 2A). These observations demonstrate that the enhanced platelet accumulation at 3500 s^-1^ is accompanied by VWF accumulation, consistent with a shear-sensitive VWF unfolding.

**Figure 2:**
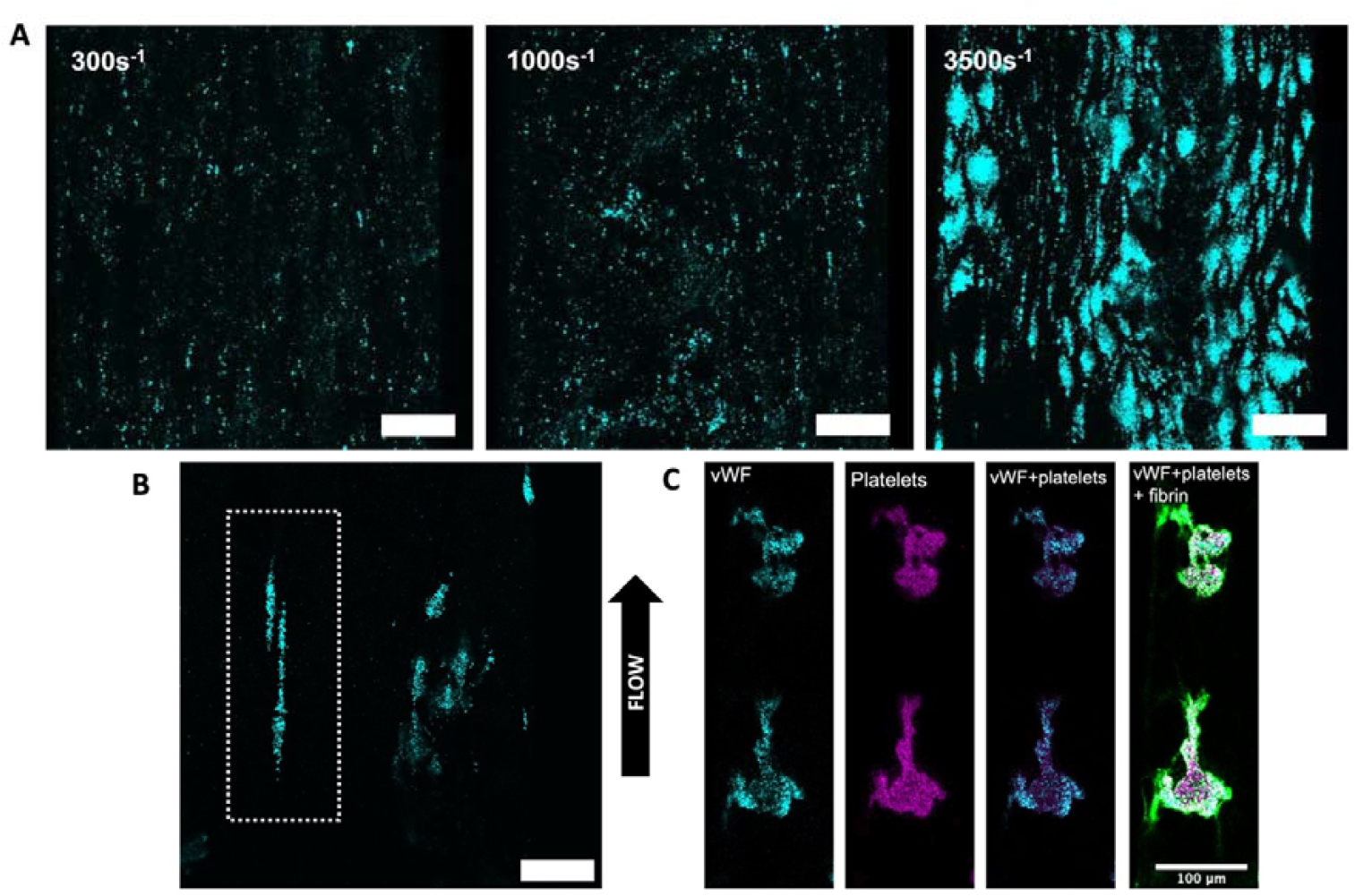
Impact of flow rate on Von Willebrand Factor (VWF) unfolding. (A) Images show VWF signal from clots formed under 300 s^-1^, 1000 s^-1^, and 3500s^-1^ at the end point of clot formation (after 10 minutes). The VWF intensity is the highest at the highest shear rate. Scale bar: 100 μm. (B) At the highest shear rate (3500s^-1^), elongated vWF strands over the collagen surface were observed (marked by dotted lines) within the first minute of flow initiation Both images show the first minute of flow, from two different experiments at 3500s^-1^. Scale bar: 100 μm. (C) Fluorescent images of VWF (cyan), platelets (magenta), merged image of vWF and platelets, and a final merged image of VWF, platelets, and fibrin (green) at 3500s^-1^ after 10 minutes. VWF and platelets colocalize while fibrin surrounds the platelet aggregate. Scale bar: 100 μm

### Final Clot Composition and Structure

To characterize the final composition of clots formed at different arterial shear conditions, we quantified the surface area covered by fibrin and platelets along with the size of platelet aggregates at the final (10 minutes) time point. Representative confocal images (Fig. 3A) show that platelet deposition significantly increased as the shear rate was increased from 300 s^−1^ to 3500⍰s^−1^. The platelet aggregates were sparse, small and spatially discrete at 300⍰s^−1^, more extensive with moderate-sized aggregates at 1000⍰s^−1^, and large and interconnected at 3500⍰s^−1^. Quantification of the final clot composition in terms of the percentage of surface area covered by platelets and fibrin at each shear rate condition showed that clots were significantly richer in platelets at the highest shear rate as compared to lower shear rates. Increasing shear lead to a progressive increase in platelet surface coverage from 9.1% ± 1.4% at 300 s^-1^, to 19.8% ± 1.1% at 1000 s^-1^ and 39.2% ± 1.5% at 3500 s^-1^ (Fig. 3B).

**Figure 3:**
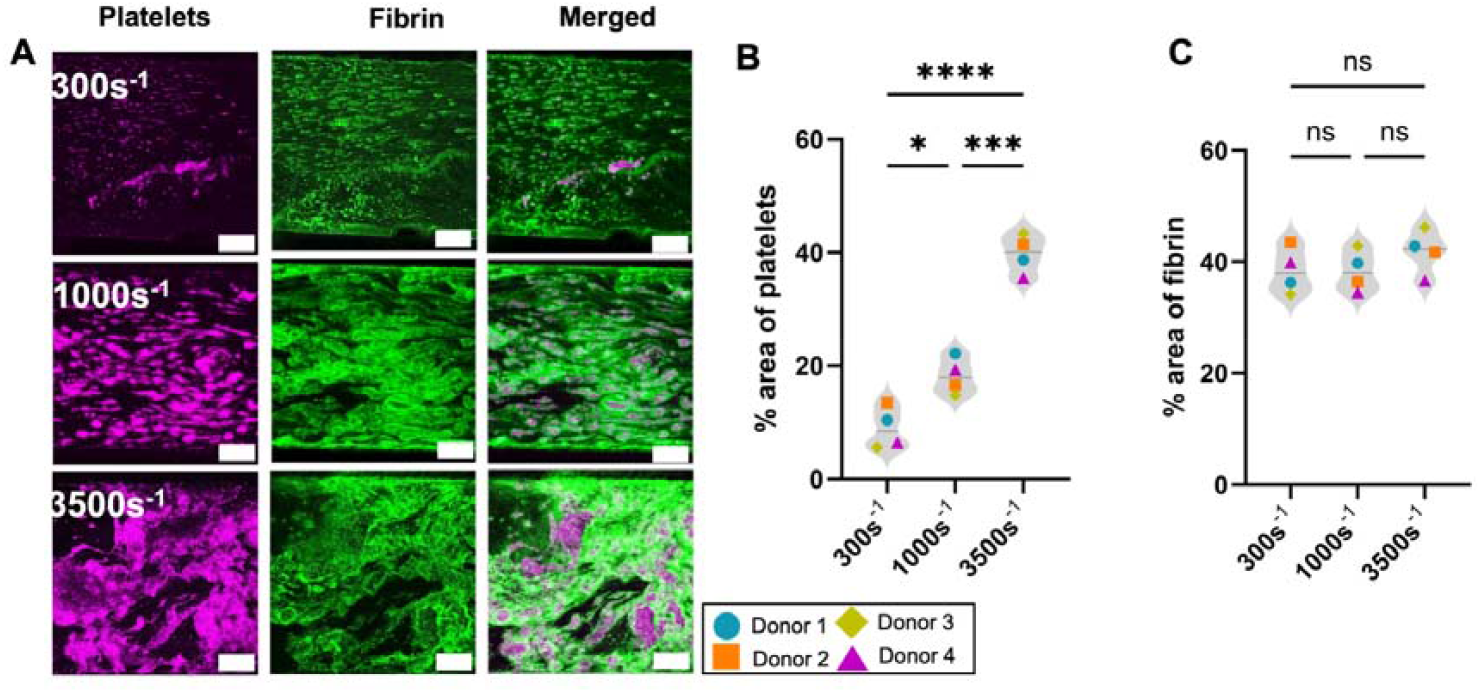
Final clot composition varies with shear rate. (A) Representative maximum intensity projections of clots formed under shear rates of 300 s^−1^, 1000 s^−1^, and 35001s^−1^ after 10 minutes of perfusion. Images display platelets (magenta) and fibrin (green). Scale bars: 100 μm. (B-C) Quantification of final clot composition in terms of the percentage of surface area covered by platelets (B) and fibrin (C) at each shear rate condition. Values for each flow rate are calculated by averaging minimum 4 replicate experiments from 4 individual donors. Higher shear rates yield clots that are significantly richer in platelets, while no clear trend is observed for fibrin coverage. (* p < 0.05, ** p < 0.005, *** p < 0.0005, **** p < 0.0001, ns: not significant)

Meanwhile, the fibrin signal showed no clear change with varying flow rates (Fig. 3A). Quantification of the fibrin surface coverage confirmed that the shear rate did not influence fibrin deposition (Fig. 3C). As a consequence, clots formed at the highest shear rate of 3500 s^-1^ were relatively richer in platelets (ratio of fibrin-to-platelet coverage 1.0), while clots formed at lower shear rates were more fibrin-rich (ratio of fibrin-to-platelet surface coverage of 3.5 for 300 s^−1^ and 1.6 for 1000 s^−1^). Quantification for each individual donor can be found in SI Figure 4.

Comparing the fibrin and platelet signals, we clearly see that the platelets are attached to the bottom surface, and are covered by fibrin (SI Figure 5). This observation is consistent with the time-lapse images from Fig. 1, which showed that platelets first adhere and aggregate on the coated surface, acting as nucleation sites for the subsequent formation of a fibrin network.

Additionally, shear rate influenced the height of the clots. This is illustrated in Fig. 4A, which shows 3D reconstructions of the clots that are color coded for depth to visualize the clot height (additional 3D renderings in Supplemental Movie 3). The average height of the clots increased with increasing shear rate, from 29 ± 1 μm at 300 s^-1^, to 40 ± 1 μm at 1000 s^-1^, and 49 ± 1 μm at 3500s^-1^ (Fig. 4B). This height increase implies an increased occlusive capacity: clots formed under 300 s^−1^ reached 41% of the channel height, while at 1000 s^−1^ they reached 57% and finally, at 3500s^−1^, they reached 70% of the channel height. Together with the increased clot height at increasing shear rate, the lateral platelet aggregate size increased significantly (Fig. 4C). At 3500 s^-1^, the largest platelet aggregates had a mean size of 3120 ± 115 μm^2^. Moreover, the aggregates were interconnected to form larger networks. The average platelet aggregate sizes at 1000 s^−1^ and 300 s^−1^ shear rates were significantly smaller, measuring 578 ± 47 μm^2^ and 67 ± 6 μm^2^, respectively.

Altogether, our data show that higher arterial shear rates enhance platelet recruitment and promote the formation of larger and taller platelet aggregates, thus enhancing the clot’s occlusive capacity.

**Figure 4:**
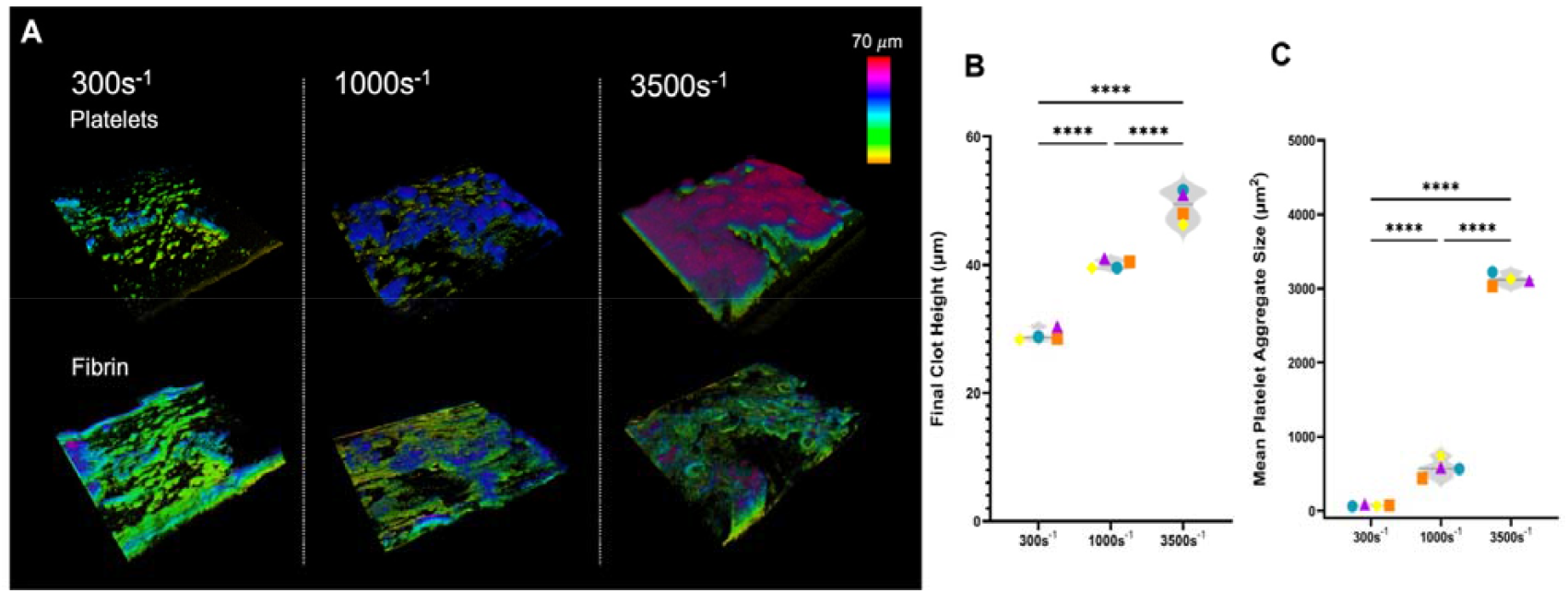
Final clot height and lateral platelet aggregate size vary with shear rate. (A) Representative 3D reconstruction of the final platelet aggregates (top row) and fibrin (bottom row) of clots formed under 300 s^-1^, 1000 s^-1^ and 3500 s^-1^. The 3D reconstructions are color coded for depth to better visualize clot height. Note that the fibrin network covers the platelet aggregates. (B) Clot height at varying shear rates, measured from the confocal z-stacks at the final (10 min) time point. At higher shear rates, clots grow taller in height (each symbol represents a donor, averaged from minimum 4 replicates per donor). (C) Quantification of the lateral size of the platelet aggregates formed under varying arterial shear from maximum intensity projections (each symbol represents a donor, averaged from minimum 4 replicates per donor). Significantly larger platelet aggregates form at higher shear rates, especially for the highest shear rate (3500 s^-1^). (**** represents p < 0.0001)

### Mechanical Properties of Clots

To determine the mechanical properties of the clots, we performed microindentation measurements on the clots using an indenter mounted on an inverted microscope. We studied clots formed at the lowest (300⍰s^−1^) and the highest (3500⍰s^−1^) shear rates, for which confocal imaging revealed large compositional differences. Fluorescence imaging showed heterogeneous distributions of platelets and fibrin at both shear rates. At the low shear rate of 300 s^-1^, we observed fibrin-rich clots and small discrete platelet aggregates (Fig. 5A). In contrast, at the high shear rate of 3500 s^-1^, we observed platelet-rich clots with large platelet aggregates surrounded by fibrin (Fig. 5B). These differences are consistent with the confocal images shown before. Simultaneous brightfield imaging revealed the presence of RBCs within the clot that did not always colocalize with fibrin or platelets, indicating the presence of regions rich in RBCs only. Quantification of the relative RBC surface coverage of these clots is presented in the supplemental materials (SI Figure 6). At both shear rates, the clots were spatially heterogeneous, showing distinct areas that could be classified as RBC-rich (Fig. 5D), RBC- and fibrin-rich (Fig. 5C) or fibrin- and platelet-rich (Fig. 5D).

To quantify the impact of the shear rate during clot formation on clot stiffness, we performed 5 indentations per clot and compared the average effective Young’s modulus (E_eff_). On average, clots formed under high shear (3500⍰s^−1^) had a significantly higher E_eff_ of 14.1 ± 1.4 kPa compared to those formed at low shear (300⍰s^−1^), with 4.1 ± 0.6 kPa (Fig. 5E). However, since clots in both cases exhibited local compositional heterogeneity, we next evaluated the mechanical heterogeneity within the clots according to the three primary groups identified from the fluorescence and brightfield images (Fig. 5C, D). As shown in Fig. 5F, irrespective of shear rate, platelet and fibrin-rich regions were significantly stiffer (17.5 ± 1.1 kPa) than areas dominated by fibrin and RBCs (6.8 ± 0.6 kPa). Regions rich in RBCs alone showed the lowest average E_eff_ of 3.1 ± 0.3 kPa, underscoring the key mechanical contribution of platelet aggregates. In conclusion, the higher average stiffness of clots formed under high shear (3500⍰s^−1^) compared to those formed at low shear (300⍰s^−1^) can be explained by the prevalence of areas rich in large platelet aggregates formed under higher flow.

## Discussion

In this study, we developed a microfluidic arterial thrombosis model to investigate how varying arterial shear rates influence clot composition, clot formation dynamics, and local mechanical properties. By combining real-time fluorescent imaging with post-formation microindentation, we provide an integrated view of thrombus development and structure along with mechanical heterogeneity under physiologically and pathologically relevant flow conditions.

### Mechanism of clot formation and final clot characteristics are shear dependent

Time lapse imaging of the clotting process showed a clear sequence of events, as summarized in Figure 6. At physiological shear rates (300 s^−1^ and 1000 s^−1^), clotting followed three main steps: (1) platelet adhesion on the coated surface, (2) platelet aggregation into discrete “islands” that expand across the surface, and (3) fibrin network formation that surrounds and interconnects aggregates, reinforcing the structure. At 300 s^−1^, overall platelet coverage remained low across 10 min and fibrin formed around relatively small aggregates. At 1000 s^−1^, platelet coverage was increased and reached a plateau with fibrin network formation. RBC retention followed the initial platelet aggregation, as the RBCs colocolized with platelet aggregates. Fibrin network formation then stabilized both the platelets and the entrapped RBCs, while streams of flowing RBCs could still move in between these aggregates, represented in supplementary movies 4-6.

**Figure 5:**
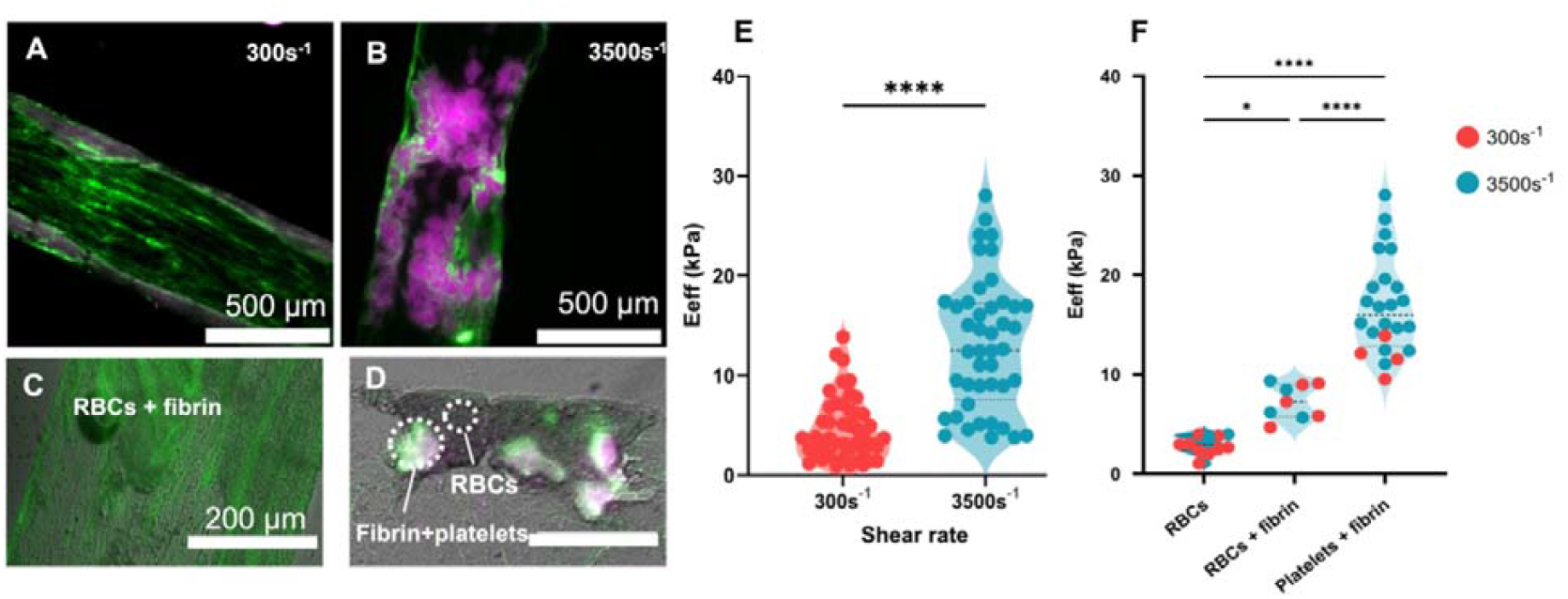
High shear rate clots are stiffer than low shear rate clots on average, but the clots show composition-dependent spatial variations in local stiffness in both cases. (A) Representative low (300 s^-1^) shear rate clot as imaged during indentation via a widefield fluorescent microscope (platelets: magenta, fibrin: green). Scale bar: 500 μm. (B) Corresponding example of a high (3500 s^-1^) shear rate clot. Scale bar: 500 μm. (C) Example of a local region rich in red blood cells (brightfield) and fibrin within a low shear rate clot (green; fluorescent channel). Scale bar: 200μm. (D) Example of a region rich in red blood cells alone versus a neighboring region rich in both platelets and fibrin within a high shear rate clot (labeled with white text). Red blood cells (RBCs) shown in brightfield, fibrin in green and platelets in magenta. Scale bar: 200 μm. (E) Effective Young’s modulus (E_eff_) of clots at low (300s^-1^) vs. high (3500s^-1^) shear rates, averaged over 9 clots measured at 5 locations. (F) Effective Young’s modulus values pooled for the two shear rates, separated by local clot composition. Regions rich in fibrin and platelets are stiffer than regions that are poor in platelets but rich in red blood cells (with or without fibrin). Regions that are rich in only red blood cells are the softest. (*p < 0.05, **** p < 0.0001)

**Figure 6:**
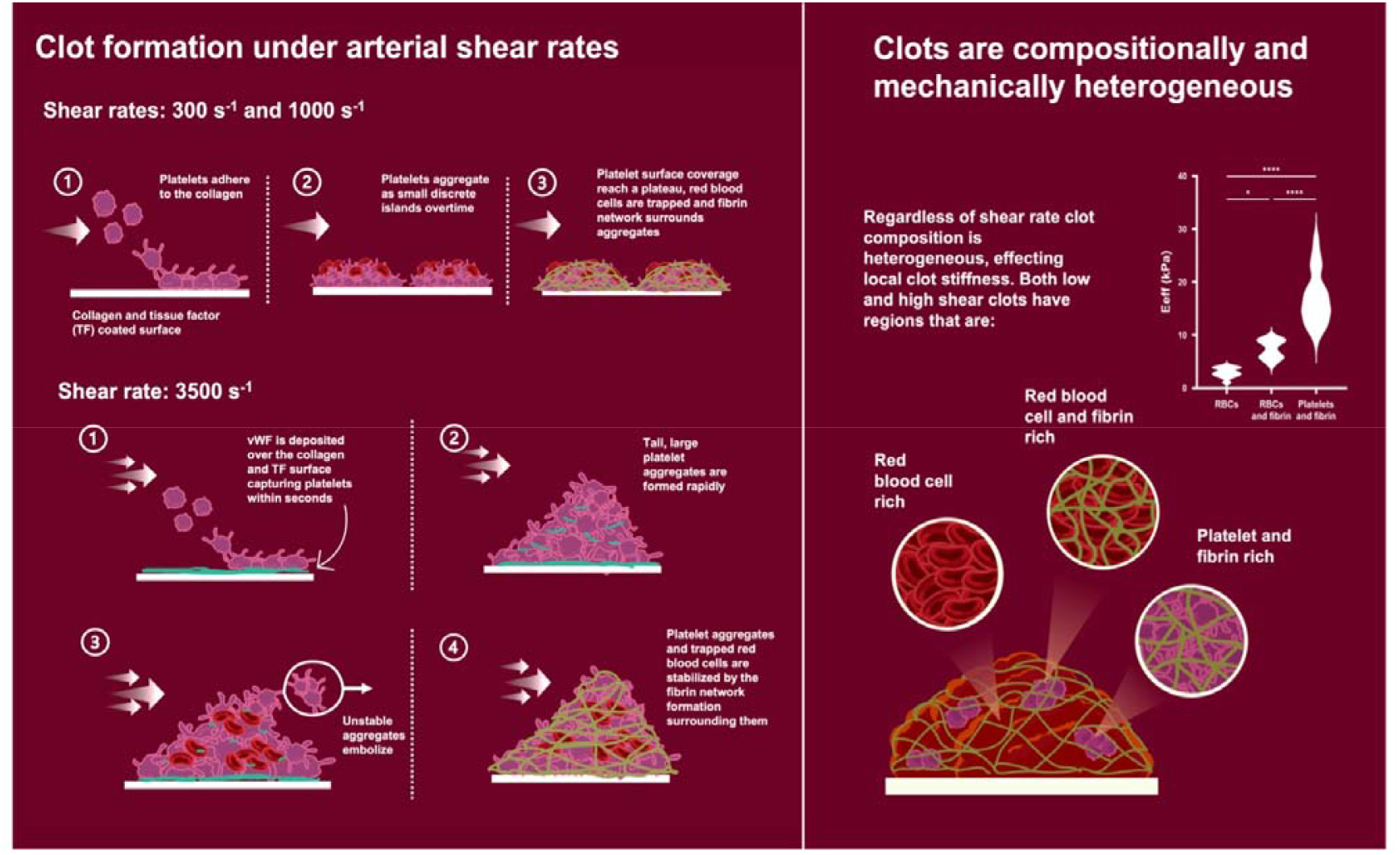
Summary of the clot formation mechanism observed under physiological (300 s^-1^ and 1000 s^-1^) and pathological (3500 s^-1^) shear rates. At 300 s^-1^ and 1000 s^-1^ the clot formation: **(1)** initiates by platelet adhesion over the coated surface, **(2)** followed by aggregation of platelets into ‘islands’ that begin entrapping red blood cells; the aggregate size increases with increasing shear rate and reaches a plateau when **(3)** fibrin network forms around the aggregates. **At 3500 s**^**-1**^, **where the shear rate is high enough for unfurling VWF: (1)** VWF strands are deposited over the coated surface facilitating platelet capture and adhesion, **(2)** tall, large platelet aggregates rapidly form, **(3)** unstable platelet aggregates embolize under high shear forces **(4)** finally they are stabilized by the fibrin network. **Regardless of the shear rate, clot composition show local heterogeneity, affecting its local stiffness**. On average, high shear rate clots are significantly stiffer than low shear rate ones. Both high and low shear rate clots have regions that are red blood cell-rich, red blood cell- and fibrin-rich, and platelet-and fibrin-rich, that show the lowest and the highest local stiffness responses, respectively.

At pathologically high shear rates (3500 s^−1^), these same steps occurred, but the initiation and growth phases were accelerated and driven by VWF. Based on previous studies of the mechanosensitivity of VWF [29, 30], 3500s^-1^ is above the shear rate threshold for VWF unfurling. At 3500 s^-1^, already within the first minute of flow, we observed VWF strands in the direction of flow, deposited over the coated surface. These strands were absent at lower rates. VWF is known to facilitate platelet capture over thrombogenic surfaces, like collagen, at high shear rates [31, 32]. We observed these strands mainly at the bottom of the coated channel surface, indicating that they form an initial bridge between collagen and platelets. This is further strengthened by the observation that the platelets and VWF colocalize in these clots. Additionally, clots formed at 3500 s^−1^ exhibited a pronounced increase in platelet aggregation and VWF signal compared to clots formed at the two lower shear rates. This can be explained by the ability of VWF to facilitate platelet-platelet aggregation at higher shear rates by tethering circulating platelets to the already adhered platelets [12, 31]. Following the rapid platelet aggregation step, an additional step emerged at this high shear: a transient unstable/embolization phase. During this unstable phase, adhered platelets detached and were carried away by the flow, causing reorganization of the platelet aggregates. This phase likely reflects the mechanical challenge faced by rapidly forming, tall platelet aggregates exposed to high shear stresses. Clots formed at 300 s^−1^ and 1000 s^−1^ are shorter and composed of smaller platelet aggregates, consistent with Pero et al. [17], who reported increased platelet surface coverage and clot height when shear was raised from 500 to 1000 s^−1^. The larger clot height at 3500 s^−1^ likely amplifies local shear rate, reinforcing platelet recruitment, whereas the shorter structures at lower shear experience less disruptive stress. This observation aligns with classical shear-induced platelet aggregate (SIPA) mechanisms described by Nesbitt et al. [11], who observed cycles of platelet aggregation and disaggregation in an *in vivo* mouse stenosis model [11]. Similarly, Colace et al. [33] showed a flow model where they first formed an initial stable platelet layer over a collagen surface at 200s^-1^, and then increased the flow rate to above 7800s^-1^, triggering a rapid (within 100 s) and unstable platelet aggregation and embolization with a simultaneous increase in VWF signal. In the same system, inhibition of GPIbα or αIIbβ3 abolished this unstable rapid aggregation and embolization burst. In our system, this unstable phase resolved when fibrin network matured (∼6 min) and stabilized platelet coverage. Our data strongly suggest that the formation of a fibrin scaffold plays a crucial role in stabilizing aggregates and halting further dynamic remodeling of the clot. These findings are consistent with prior work demonstrating that fibrin cross-links and consolidates platelets, resisting disintegration and removal by flow [14, 34, 35].

### Clot composition determines local stiffness, with platelet-rich regions being stiffer

Indentation measurements showed that clots formed under high shear (3500⍰s^−1^) were on average 3.4-fold stiffer than clots formed at low shear (300⍰s^−1^). This finding is consistent with an earlier study, where clots formed in a stenotic microfluidic model were 2.4-fold stiffer when formed under high shear (3500s^-1^) as compared to low shear flow [19]. We observed marked regional mechanical heterogeneity for clots across all shear conditions. This mechanical heterogeneity co-varied with local clot composition: regions that were rich both in platelets and fibrin were significantly stiffer than regions dominated by fibrin and RBCs, while regions mainly rich in RBCs were softest. Similar heterogeneity was observed in thrombi retrieved from patients [15, 36]. We therefore attribute the larger stiffness of clots formed at higher shear to their larger platelet content and platelet-to-fibrin ratio, clearly observed by imaging. Although we did not measure contraction directly, platelet-driven clot contraction is known to compact fibrin and RBCs, leading to polyhedrocyte formation, providing a plausible mechanistic link between platelet content and increased stiffness [37-39]. Indentation measurements by Swieringa et al. on clots formed in a microfluidic device with a single channel [14] similarly revealed stiffer clots at high (1000 s^-1^) as compared to low (150 s^-1^) shear with a larger platelet-to-fibrin ratio. Our observation of lower Young’s moduli for RBC-rich regions aligns with previous macroscopic compression and tensile measurements by Cahalane et al. [18] for static clot analogues with varying RBC content, where increasing RBC concentration lowered the clot stiffness. Our observations are likewise consistent with analyses of human ischemic stroke thrombi, which reported a strong positive correlation between fibrin/platelet content and stiffness, and an inverse correlation between RBC content and stiffness [4]. Altogether, our findings confirm the key contribution of platelet aggregates to thrombus stiffness, as previously observed in *ex vivo* thrombi and *in vitro* static clots [6, 7, 18].

Traditional analyses of patient-derived clots are mainly performed on thrombi that are post-embolization, altered due to thrombectomy procedure or treatment, and furthermore it is not possible to identify the structural differences due to the time delay between analysis and thrombus formation. Here, we demonstrate an experimental platform to generate, structurally and mechanically study blood clots from the moment of formation under flow, offering a more complete and unbiased perspective of thrombus evolution. Furthermore, microindentation allows spatially resolved mechanical mapping, which is not achievable through bulk testing. This approach may offer new insight into which regions of a thrombus are most vulnerable to embolization or most resistant to thrombectomy.

### Limitations

While our model replicates arterial shear conditions and offers high-resolution imaging, there are several aspects where it differs from the *in vivo* situation: it does not include an endothelial cell layer nor pulsatile or gradient flow, which can modulate platelet adhesion and fibrin formation *in vivo* [40, 41]. All experiments were conducted at room temperature which may underestimate the amount of platelet adhesion, aggregation and contraction [42]. Similarly, the rectangular cross-section of the channel does not fully recapture the *in vivo* circular vessels, leading to a nonuniform distribution of wall shear stress towards the corners of the channel [43]. The microindentation analysis assumes Hertzian contact with a homogeneous, purely elastic material; given the viscoelastic and porous nature of clots [44, 45], the reported E_eff_ values should be interpreted in relative rather than absolute terms. RBC-rich regions were classified using brightfield imaging; more quantitative staining could refine the assessment of RBC content.

## Conclusions

We studied clot formation across a range of arterial shear rates, revealing how, within the arterial regime, shear rate influences clot structure, growth dynamics, and mechanical properties. Our findings reveal that increasing shear rates promote faster formation of larger platelet aggregates, but also introduce a transient instability phase prior to fibrin stabilization at pathologically high shear rates. We further demonstrate that platelet- and fibrin-rich regions are significantly stiffer than RBC-dominant regions, underscoring how compositional heterogeneity translates into mechanical heterogeneity at the microscale. By enabling observation and analysis of the full course of clot development under physiologically relevant conditions, our platform bridges an important gap by allowing local mechanical characterization coupled with compositional information to account for heterogeneity of thrombi.

## Supporting information

SI mov 6

SI mov 5

SI mov 3

SI mov 4

SI mov 1

SI mov 2

## Acknowledgements

This project was funded by the Dutch Thrombosis Foundation Trombosestichting with grant number 2021_02. We thank Michiel Manten (Erasmus MC) for his help with designing and 3D Printing the master mold and Irène Nagle (TU Delft) and Malin Becker (Optics11) for their invaluable help with the microindentation measurements and data analysis. We also thank Iris van Moort (Erasmus MC) for providing the VWF antibody. Finally, we thank Frank Gijsen (TU Delft) for sharing his insights on experimental design and for helpful discussions.

## Author Contributions

Hande Eyisoylu conceived and designed the research, performed the experiments, analyzed the data, and wrote the manuscript. Parvin Ebrahimi performed the experiments and analyzed the data. Marla Lavrijsen contributed to the methodology and interpretation of the data. Gijsje H. Koenderink and Moniek P.M. de Maat contributed in conceptualization, supervision, writing-review and editing. All authors approved the final version of the manuscript.

