## Supplemental Figures for "Arterial Shear Rate Determines the Structure and Mechanical Properties of Whole-Blood Clots"

**Supplemental Information**

**Microfluidic Model Fabrication and Assembly:**

***Master Mold Fabrication:***

The master molds for the microfluidic devices were fabricated via 3D printing using Envisiontec VIDA-HD printer, employing Envisiontec HTM 140 as the printing material (Envisitiontec). The high resolution (50μm XY and 25μm Z resolution) and mechanical stability of this material ensure the precision and durability of the mold features required for microfluidic device fabrication.

***PDMS Device Fabrication:***

The microfluidic devices were fabricated using poly(dimethylsiloxane) (PDMS) prepared from the SYLGARD-184 silicone elastomer kit (VWR Chemicals). The PDMS prepolymer and curing agent were mixed at a ratio of 10:1 (w/w). To remove entrapped air bubbles, the mixture was degassed for an hour in a desiccator. Degassed PDMS was poured over the 3D-printed molds and allowed to settle. The PDMS-coated molds were baked overnight at 60°C to cure. After curing, the PDMS layers were carefully unmolded. The inlets and outlets were punched in at the marked locations using a biopsy punch.

***Assembling of the Flow Device:***

For reversible bonding, the PDMS mold was bonded to the coverslip using a vacuum-sealing method. The device design incorporated a “vacuum chamber” surrounding channel walls, separate from the main channel (SI Figure 1). The vacuum-sealing was administered using a 1 mL syringe, and a clamp to secure the syringe. The vacuum could be released at the end of the experiments to allow access to the formed clot. This approach ensured a secure seal between the PDMS and coverslip, enabling reliable experiments without leakage.

**Supplementary Figures**

**A**

**B**

**C**

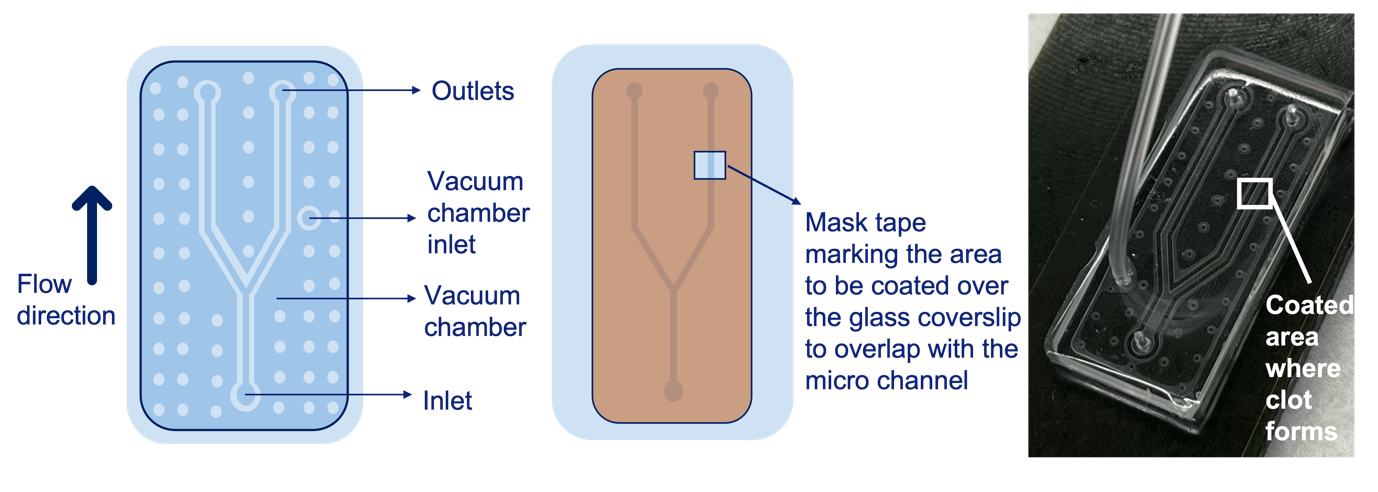

**SI Figure 1: Microfluidic flow device assembly.** A) A schematic of the flow model marking the inlets, outlets, main channels, and the surrounding vacuum chamber. The main channels are 500 μm wide and 70 μm tall. B) Representation of the mask use to mark the area to be coated with collagen and tissue factor, which will overlap with the flow channel. To make sure the coated area is the same size and at the same location for each experiment, a mask, as shown in B in orange, was placed over the glass coverslip. The collagen and tissue factor were pipetted over this spot. C) A photograph of the assembled flow model. Vacuum is applied via the tubing connected to the vacuum chamber inlet. This allows for reversible bonding that is secure during thrombus growth under varying shear rates, while allowing access to the clot by releasing the vacuum and gently lifting off the PDMS chamber. (The PDMS block measures 1.5 cm x 3.5 cm.)

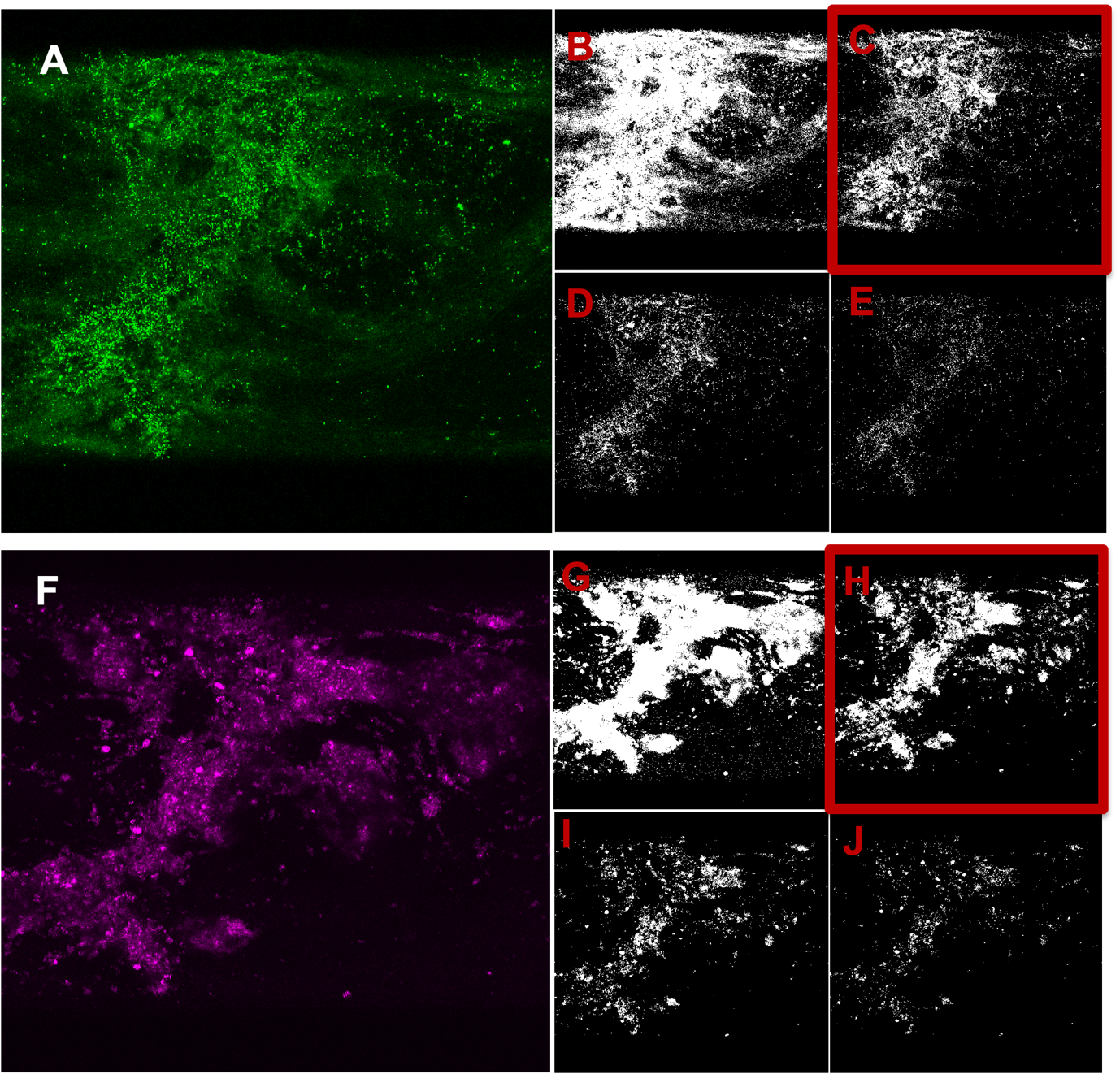

**SI Figure 2: Comparison of automatic thresholding methods for fibrin and platelet channels.**

Representative maximum projections of confocal Z-stacks of a clot formed at 1000 s^-1^ showing the fibrin (A) and platelet (F) channels, alongside binary masks generated by four automatic thresholding algorithms: Default (B and G), Otsu (C and I), Triangle (D and H) and IsoData (E and J). The selected threshold masks for each channel are marked with the red box: Otsu (C) represented the fibrin channel the best by capturing the intense fibrin regions but also including the more faint fibrin networks without capturing too much background noise. For the platelets channels, triangle (H) represented the platelet signal the best by capturing minimal noise while closely resembling the platelet aggregates in the original image.

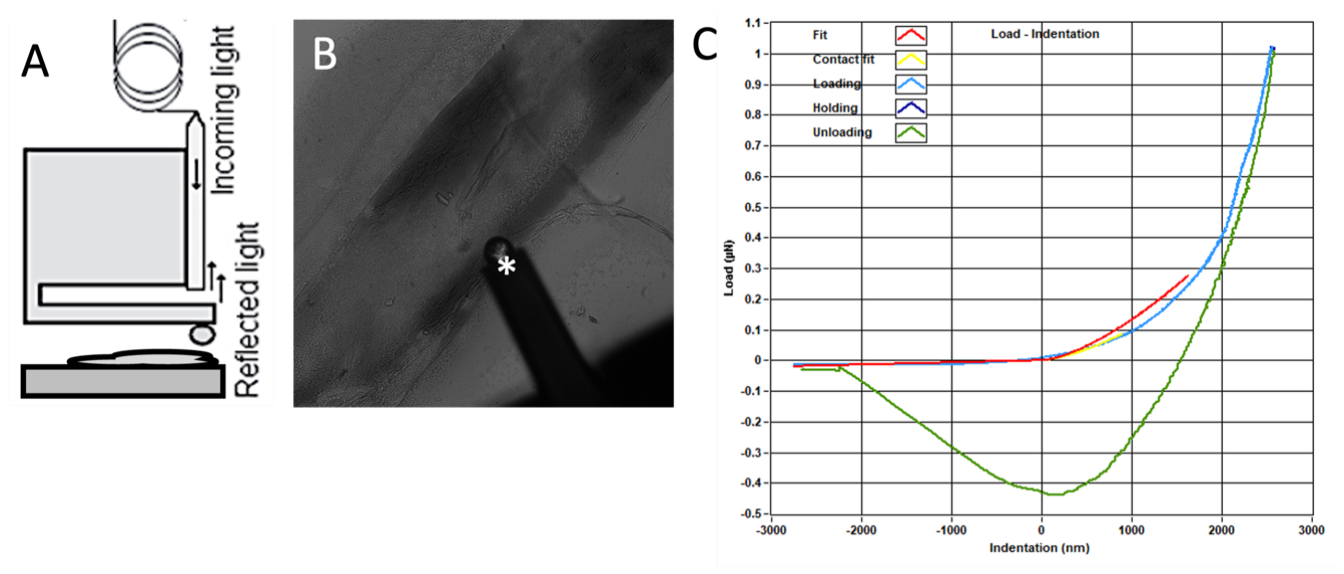

**SI Figure 3: Microindentation setup.** A) A schematic representation of the microindentation setup, which is mounted on an inverted microscope. The micro-indenter consists of a probe and an interferometer. The probe has an optical fiber and a cantilever with a spherical tip of radius 56 µm. The probe displacement and the forces applied are measured via the fiber-optic interferometry. B) Representative brightfield image showing a clot sample (which runs diagonally from bottom-left to top-right) with the microindenter probe (marked with *) positioned on top. C) A representative indentation curve showing the Hertz Contact fit (yellow) over the loading curve (blue) and the unloading curve (green) on a single indentation. By fitting this curve on the Hertz model equation (equation 2 in the main text), the effective Young’s Modulus of the sample at the location of the indentation is calculated.

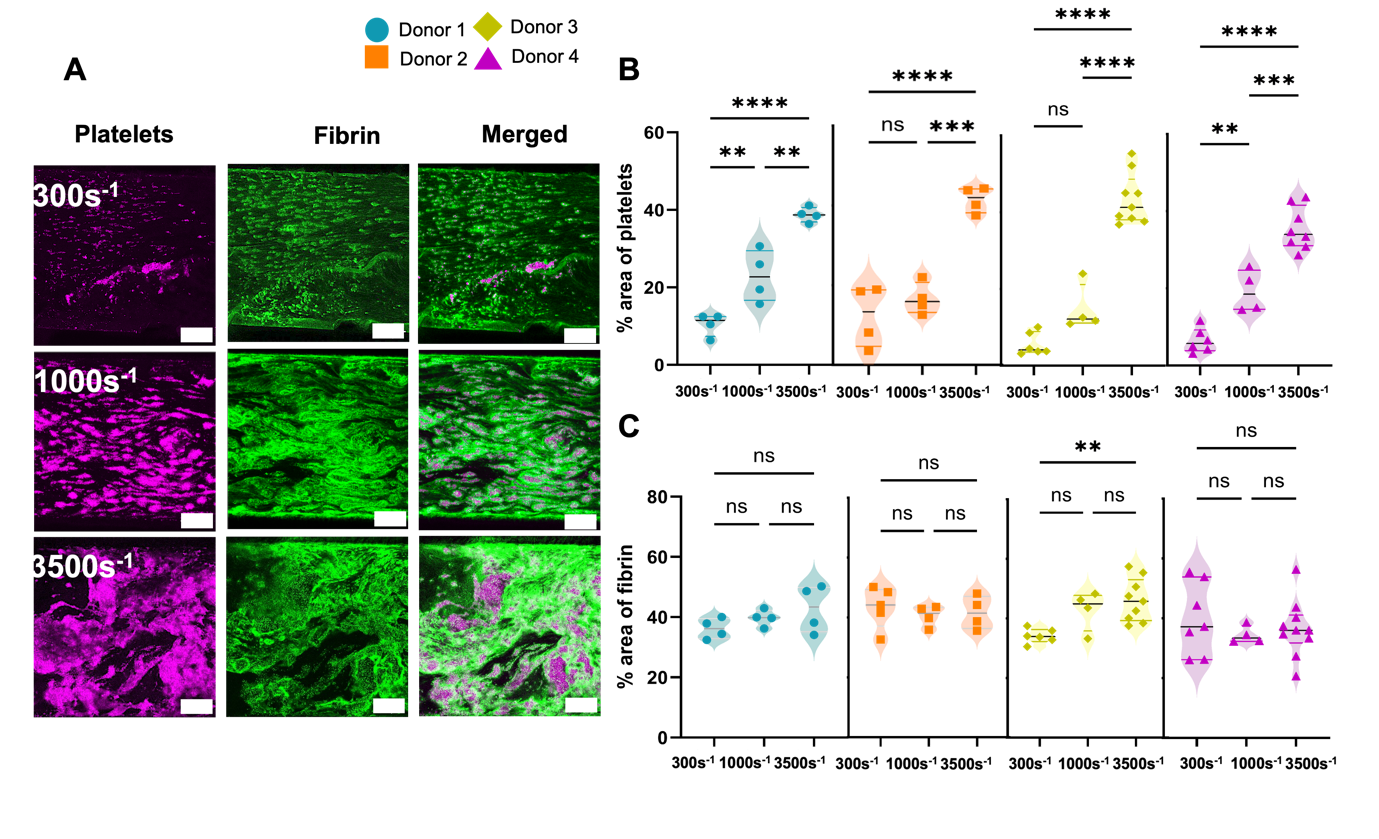

**SI Figure 4: Final clot composition varies with shear rate for each donor**. (A) Representative maximum intensity projections of clots formed under shear rates of 300 s⁻¹, 1000 s⁻¹, and 3500 s⁻¹ after 10 minutes of perfusion. Images display platelets (magenta) and fibrin (green). Scale bars: 100 μm. (B-C) Quantification of final clot composition in terms of the percentage of surface area covered by platelets (B) and fibrin (C) at each shear rate condition, separated per individual donor (color-coded). For all donors, at highest shear rate, clots were significantly richer in platelets compared to the lower shear rates, two of the donors show a significant difference in platelet coverage between the 300 s⁻¹ and 1000 s⁻¹ conditions as well. (C) No significant difference is found in fibrin coverage due to flow rate except for one donor between the higest and the lowest shear rates.

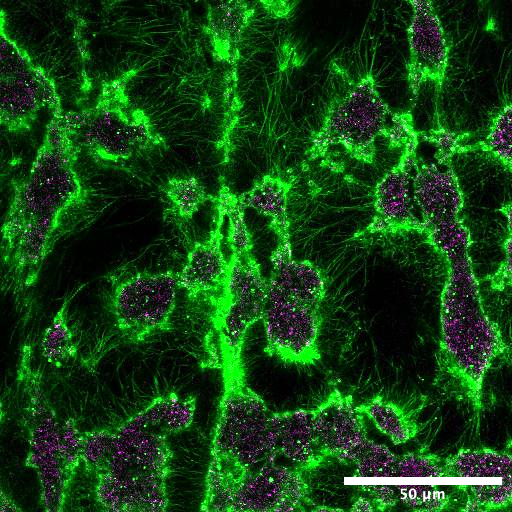

**SI Figure 5: Close-up view of platelet aggregates and fibrin distribution in a clot.** A maximum intensity projection of a confocal fluorescence Z-stack of a representative clot formed under 1000s^-1^ shear rate after 10 minutes of perfusion. Platelet aggregates (magenta) are formed as ‘islets’ over the tissue factor/collagen I-coated surface. Fibrin fibers (green) surround and interconnect the platelet aggregates. Scale bar: 50 µm.

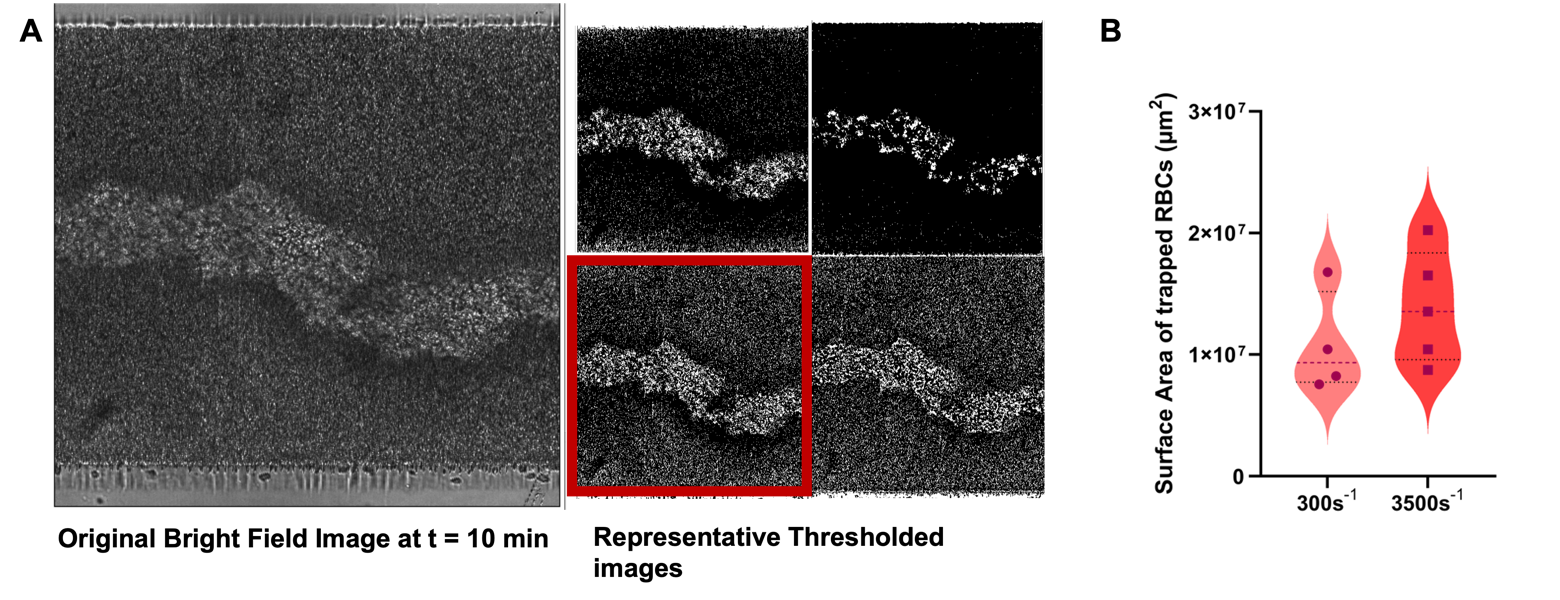

**SI Figure 6: RBC surface area quantification of the final clots at the highest (3500 s^-1^) and the lowest (300 s^-1^) shear rates.** (A) A representative brightfield image of a clot formed over 10 minutes at 3500s^-1^ is shown on the right. Brightfield (BF) images were acquired concurrently with confocal z-stacks of fibrin and platelets. Because BF lacks optical sectioning, RBCs dominate BF contrast under these conditions. The brightfield image at the final time point was taken as the fully formed clot. For quantification, we extracted a single best-focus BF plane per field. The area occupied by entrapped red blood cells (RBCs) was measured. Briefly, in ImageJ (NIH) the final brightfield image was converted to 8-bit, and segmented using a threshold chosen to most closely visually match the original image while excluding background signal (non-entrapped, still-flowing RBCs and the channel walls). The left panel show representative examples of thresholded images, highlighting the selected one with the surrounding red box. The resulting binary mask of the clot region was used to calculate the clot area (µm²). This procedure was applied to clots formed at the lowest (300 s^-1^) and the highest (3500 s^-1^) shear rates, and the measured clot areas were compared between these conditions. Additionally, we visually confirmed that the mask corresponded to “trapped” RBCs by reviewing earlier timepoints in the time-lapse where flowing versus adherent RBCs could be distinguished, and only retained regions that remained stationary across frames. BF signal integrates through the clot thickness and is not specific for RBCs; however, in our flow model at 20X, phase-dense BF features are expected to be predominantly RBCs rather than fibrin or platelets because of their abundance and size. In experiments imaged overtime, platelet fluorescence due to adhesion and aggregation typically appears before any BF-visible aggregation in the same region; we interpret the subsequent BF increase within the clot area mask as the arrival/retention of RBCs following platelet aggregation. The resulting BF-derived value is reported as a relative RBC coverage for within-assay comparisons across shear conditions; it is not interpreted as an absolute RBC count or volume fraction. (B) The surface area coverage of RBCs in the brightfield channel at the lowest (300 s^-1^) and the highest (3500 s^-1^) shear rates.
